# O-GlcNAcylation regulates PPAR-driven metabolic programming in intestinal stem cells

**DOI:** 10.64898/2026.03.13.711696

**Authors:** Thomas Hartley McDermott, Dominic R Saiz, Yesenia Barrera Millan, Ngoc Bao Phuong Ho, Matthew Torel, Eric Uher, Caleb Aboagye, Fiona Farnsworth, Gourab Lahiri, Venkataramana Thiriveedi, Jinhua Chi, Haiwei Gu, Charlie Fehl, Benjamin B Bartelle, Miyeko D Mana

## Abstract

Diet deeply influences health and disease risk by reshaping cellular metabolism. In the intestine, dietary nutrients directly affect intestinal stem cell (ISC) behavior, yet the regulatory mechanisms linking metabolism to transcriptional control remain poorly defined. Because mitochondria function as central metabolic hubs, we focused on mitochondrial signaling to understand how nutrient utilization governs ISC function. Using the MITO-Tag mouse, we isolated metabolites specifically from ISC mitochondria and found that the sugar-derived metabolite UDP-GlcNAc was reduced in ISCs from mice fed a high-fat diet. Moreover, we identified that reducing O-GlcNAcylation (OGN) rapidly increased stem cell frequency, proliferation, regenerative capacity, and the abundance of PPAR target proteins. Mechanistically, these effects depend on PPAR signaling, as genetic loss of *Ppar-d/a* blocks the ISC phenotypes induced by reduced OGN. These results reveal an OGN–PPAR signaling axis that translates dietary metabolic cues into transcriptional programs governing fuel utilization and ISC behavior in the intestine. Collectively, our findings highlight that OGN is a previously unrecognized regulator of PPAR signaling in intestinal stem cells.

## INTRODUCTION

Adult tissue stem cells preserve tissue integrity by coupling metabolic state to transcriptional programs that influence proliferation, self-renewal, and differentiation (Jackson and Finley, 2024; Ly et al., 2020; Meacham et al., 2022; Shapira and Christofk, 2020). Nutrient-sensing signaling pathways integrate fluctuations in nutrient input, fuel availability, and intermediary metabolites, thereby reshaping gene expression networks and determining stem cell functional output (Ghosh-Choudhary et al., 2020; Mihaylova et al., 2014; Novak et al., 2021). Through this metabolic–transcriptional integration, stem cells dynamically tune self-renewal and proliferative capacity in response to physiological demands, thus sustaining long-term tissue homeostasis while preserving regenerative potential. This axis is increasingly appreciated as a determinant of long-term tissue health, as changes in stem cell–intrinsic metabolism can profoundly impact tissue fitness and promote disease states, including cancer (Shay and Yilmaz, 2025; Burclaff, 2023; Novak et al., 2021).

The intestinal epithelium is uniquely exposed to dynamic nutrient environments, requiring continuous integration of dietary signals to coordinate metabolism. At the apex of cellular hierarchy, *Lgr5*^+^ intestinal stem cells (ISCs) continuously regenerate the epithelium while adapting to fluctuations in nutrient status (Barker et al., 2007; Beyaz et al., 2016; Cheng et al., 2019; Goncalves et al., 2019; Imada et al., 2024; C. Li et al., 2023; Mana et al., 2021; Mihaylova et al., 2018; Yilmaz et al., 2012). Mitochondria sit at the center of the metabolic–transcriptional axis because they convert nutrient inputs into energy production, biosynthetic substrates, and signaling factors that directly influence gene regulation and cell state (Wanet et al., 2015; Rath et al., 2018a; Ludikhuize et al., 2020; Martínez-Reyes and Chandel, 2020; Wang et al., 2024; Chaves-Perez et al., 2025). For example, ketogenic diets induce mitochondrial HMGCS2-mediated β-hydroxybutyrate (βOHB) production to regulate nuclear Notch signaling and enhance ISC self-renewal (Cheng et al., 2019). While high-fat diets (HFD) and fasted states augment ISC function by converging on a PPAR nuclear receptor program and the downstream transcriptional target *Cpt1a* to control mitochondrial oxidation of long-chain fatty acids (Beyaz et al., 2016; Mana et al., 2021; Mihaylova et al., 2018). These studies underscore how mitochondrial metabolism serves as a central hub through which dietary signals shape ISC adaptive responses.

To further refine our understanding of the metabolic-transcriptional axis, we asked how mitochondrial metabolite composition changes under different nutrient conditions and whether these metabolites directly regulate stem-cell function. Using MITO-tag isolation of ISC mitochondria from mice in various dietary conditions, we performed comparative metabolomics. This analysis identified the hexose amine biosynthesis pathway and its generation of uridine-diphosphate-N-acetylglucosamine (UDP-GlcNAc), linking ISC metabolism to protein glycosylation through O-GlcNAcylation (OGN), a reversible post-translational modification catalyzed by O-GlcNAc transferase (OGT)(Bayraktar et al., 2019; Chen et al., 2016). OGN has emerged as a metabolic signaling mechanism capable of regulating metabolic enzymes, transcription factors, and chromatin-associated proteins in response to nutrient flux (Love et al., 2003; Ma et al., 2015; Mannino and Hart, 2022; Ranuncolo et al., 2012; Yang and Qian, 2017). Although OGN is known to respond to glucose availability and metabolic stress, its role in coordinating lipid metabolism and transcriptional regulation in adult stem cells remains largely unexplored (Gonzalez-Rellan et al., 2022; Lockridge and Hanover, 2022; Shin et al., 2023). We show that diet-dependent mitochondrial metabolism alters UDP-GlcNAc abundance in ISCs, and that reduced OGN promotes a PPAR-driven lipid metabolic program that enhances stem-cell function. These findings identify OGN as a metabolic signaling link combining glucose utilization, lipid metabolism, and transcriptional control of intestinal stem cells, thereby defining an unexpected regulatory axis that governs how ISCs dynamically shift between glucose and lipid metabolism as competing fuel sources.

## RESULTS

### Mitochondria from HFD-induced ISCs have decreased UDP-GlcNAc

To explore the metabolic-transcriptional axis governing ISCs, we generated *Rosa26*^LSL-3XHA-EGFP-OMP25/+^; *Lgr5*^CreERT2^ mice (MITO-Tag^ISC^) to selectively mark mitochondria with an HA-epitope tag for rapid isolation via immunoprecipitation and metabolite analysis (Figure 1A). To identify the timepoint at which induced MITO-tag expression is restricted to ISCs rather than their progeny, we administered tamoxifen at variable timepoints using *Rosa26*^tdTomato^; *Lgr5*^CreERT2^ as a proxy and assessed tdTomato^+^ cells within the crypt. We identified a 40-hour timepoint with tdTomato production limited to ISCs, whereas a 60-hour induction resulted in tdTomato^+^ labelling within differentiated cells migrating higher up the crypt axis. We verified that HA-MITO expression from a 40-hour induction of MITO-Tag^ISC^ was restricted to the base of the crypt and colocalized with mitochondria using immunofluorescence (IF). No HA-MITO was detectable in the uninduced mice (Figures 1B and S1A-C).

**Figure 1.**
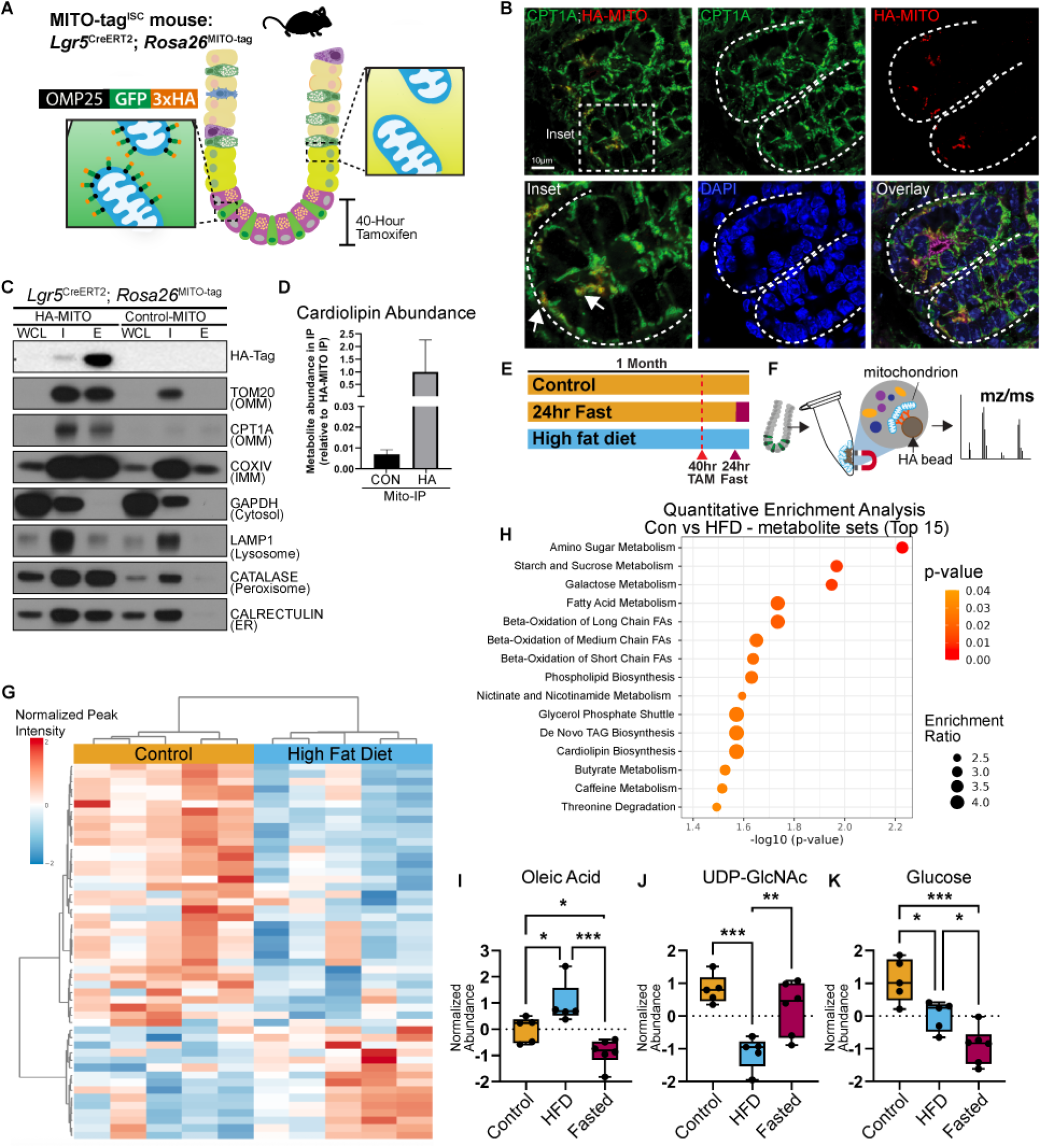
MITO-Tag reveals decreased UDP-GlcNAc in HFD Mitochondria. (A) Schematic of MITO-Tag^ISC^ Mouse. (B) Representative Airyscan images of MITO-Tag^ISC^ mouse after 40-hour induction with 100mg/kg Tamoxifen. Red: HA-MITO; Green: CPT1a; Blue: DAPI. (C) Immunoblot analysis of jejunum tissue (WCL: whole cell lysate), mitochondria enriched input (I), and anti-HA immunoprecipitated extract (E) from either HA-MITO or Control-MITO. Protein markers and their corresponding subcellular compartment are to the right. OMM = Outer Mitochondrial Membrane; IMM = Inner Mitochondrial Membrane; ER = Endoplasmic Reticulum. (D) Lipidomic analysis of mitochondrial immunoprecipitate for cardiolipin abundance in Control-MITO (N=5) and HA-MITO (N=5). (E) Schematic of MITO-Tag^ISC^ mice on Control (1 month), HFD (1 month), and Control (1 month) plus a 24 hour fast. (F) Schematic of isolation of HA-Mitochondria from ISCs for metabolomic analysis. (G) Heat map of mitochondrial metabolites between Control and HFD isolated mitochondria (N=5) (H) Quantitative enrichment analysis of mitochondrial metabolites from Control and HFD MITO-Tag^ISC^ mice. (I,J,K) Normalized abundance of oleic acid (I), UDP-GlcNAc (J) and Glucose (K) from Control, HFD and Fasted MITO-Tag^ISC^ mice. \**p<0*.*05, **p<0*.*01, ***p<0*.*001* by one-way ANOVA.

As validation of the functional isolation of HA-tagged mitochondria, immunoblot analysis showed enrichment for mitochondrial outer (TOM20, CPT1A) and inner (COXIV) membrane proteins relative to cytoplasmic proteins (GAPDH)(Figure 1C). Signal from other compartments was detected as expected given the known contacts made between mitochondria and peroxisomes (CATALASE), endoplasmic reticulum (CALRETICULIN), and lysosomes (LAMP1), consistent with previous reports (Bayraktar et al., 2019; Lewis et al., 2016; Schrader et al., 2013). Moreover, by transmission electron microscopy, we frequently observed contact sites between mitochondria, endoplasmic reticulum, and lipid droplets in ISCs (Figure S1D). Importantly, uninduced mice exhibit minimal non-mitochondrial protein contamination in the immunoprecipitate (IP)(Figure 1C). Using lipidomics, we found that the mitochondrial-specific lipid, cardiolipin, is significantly enriched in the HA-MITO IP relative to background (Figure 1D). Collectively, these data demonstrate that mitochondria can be rapidly and successfully isolated from ISCs in a manner consistent with prior reports in other tissues (Bayraktar et al., 2019; Sprenger et al., 2025).

We leveraged three dietary strategies as an entry point to test how metabolic signaling interfaces with transcriptional regulation in ISCs: a 1-month Control diet, a 1-month HFD, and, for contrast, a 24 hour fast (following 1 month on Control diet) (Figures 1E). After confirming that HA-MITO expression in OLFM4^+^ ISCs was similar between conditions (Figure S1C), we placed MITO-Tag^ISC^ mice on the various diets and performed targeted metabolomics on isolated mitochondria (Figure 1F). Using Metaboanalyst (Pang et al., 2024), we observed strong unbiased clustering of mitochondrial metabolites between control and HFD conditions, confirming that mitochondrial metabolism has distinct features in each diet (Figure 1G). For example, we detected increases in oleic acid in HFD mitochondria, likely due to increased long chain fatty acid transport for fatty acid oxidation (Figure S1E). Conversely betaine levels were decreased, consistent with HFDs decreasing betaine abundance (Figure S1E) (Beyaz et al., 2016; Ejaz et al., 2016; Mana et al., 2021). To identify the metabolic pathways altered by diet, we performed Quantitative Enrichment Analysis and observed an enrichment of Amino Sugar Metabolism (Figure 1H). In mitochondria isolated from HFD ISCs, Amino Sugar Metabolism was consistently reduced, driven in part by the differential abundance of UDP-GlcNAc, a key downstream product of hexosamine biosynthetic pathway (HBP) (Figure 1J and S1E-F). Because the HBP pathway branches from glucose metabolism, and glucose abundance was reduced under HFD conditions, this initially suggested that decreased UDP-GlcNAc might simply reflect reduced glucose availability (Figure 1K) (Paneque et al., 2023). However, comparison of control to fasted mitochondria, which also exhibit reduced glucose abundance, revealed no significant change in UDP-GlcNAc levels, indicating that the reduction observed in HFD mitochondria reflects a functional metabolic shift rather than a substrate limitation (Figure 1K). These findings link glucose and lipid metabolism through UDP-GlcNAc raising the possibility that the HBP and UDP-GlcNAc abundance serve as a regulatory nodes in ISC function.

### Reduced OGN increases stemness

The HBP diverts a fraction of glycolytic intermediates to produce UDP-GlcNAc, the essential substrate for OGN and N-linked glycosylation. OGN of intracellular proteins at serine and threonine residues by OGT regulates fundamental cellular processes, including transcription, protein stability, metabolism and cellular stress survival, but the molecular distinction in ISCs is unknown (Chu et al., 2014; Jackson and Tjian, 1988; Love et al., 2003; Ma et al., 2015; Ranuncolo et al., 2012). Elevating glucose availability in intestinal organoid cultures increased the levels of cellular OGN in a concentration-dependent manner (Figure S2A), indicating that increased glucose flux promotes OGN in intestinal epithelia. To understand the extent to which decreased UDP-GlcNAc in HFD mitochondria influences ISCs, we first utilized pharmacological inhibition of OGT (ST045849) to partially suppress OGN without fully blocking OGT activity (Figure 2A)(Itkonen et al., 2016; Zeng et al., 2025; Zhao et al., 2018). We established a working concentration (20μM) that lowers OGN levels after a 3 day administration on freshly harvested crypts while maintaining organoid formation and viability, thereby avoiding the detrimental outcomes associated with OGT loss (Figures 2B and S2B) (Xiong et al., 2022; Zhao et al., 2020, 2018). We then exposed organoids harboring an ISC GFP reporter (*Lgr5*^eGFP-IRES-CreERT2^; *Vil*^CreER^) to partial OGT inhibition for 3 days and evaluated GFP^+^ ISC frequency using flow cytometry in which decreased OGT activity resulted in increased *Lgr5*-eGFP^hi^ stem cells compared to vehicle-treated controls (Figure 2C). To assess proliferation, we measured BrdU incorporation in ST-treated *Lgr5*-eGFP^+^ organoids, and quantified BrdU^+^; GFP^hi^ ISCs via IF (Figure 2D) revealing increased proliferation in ST-treated organoids over vehicle controls.

**Figure 2.**
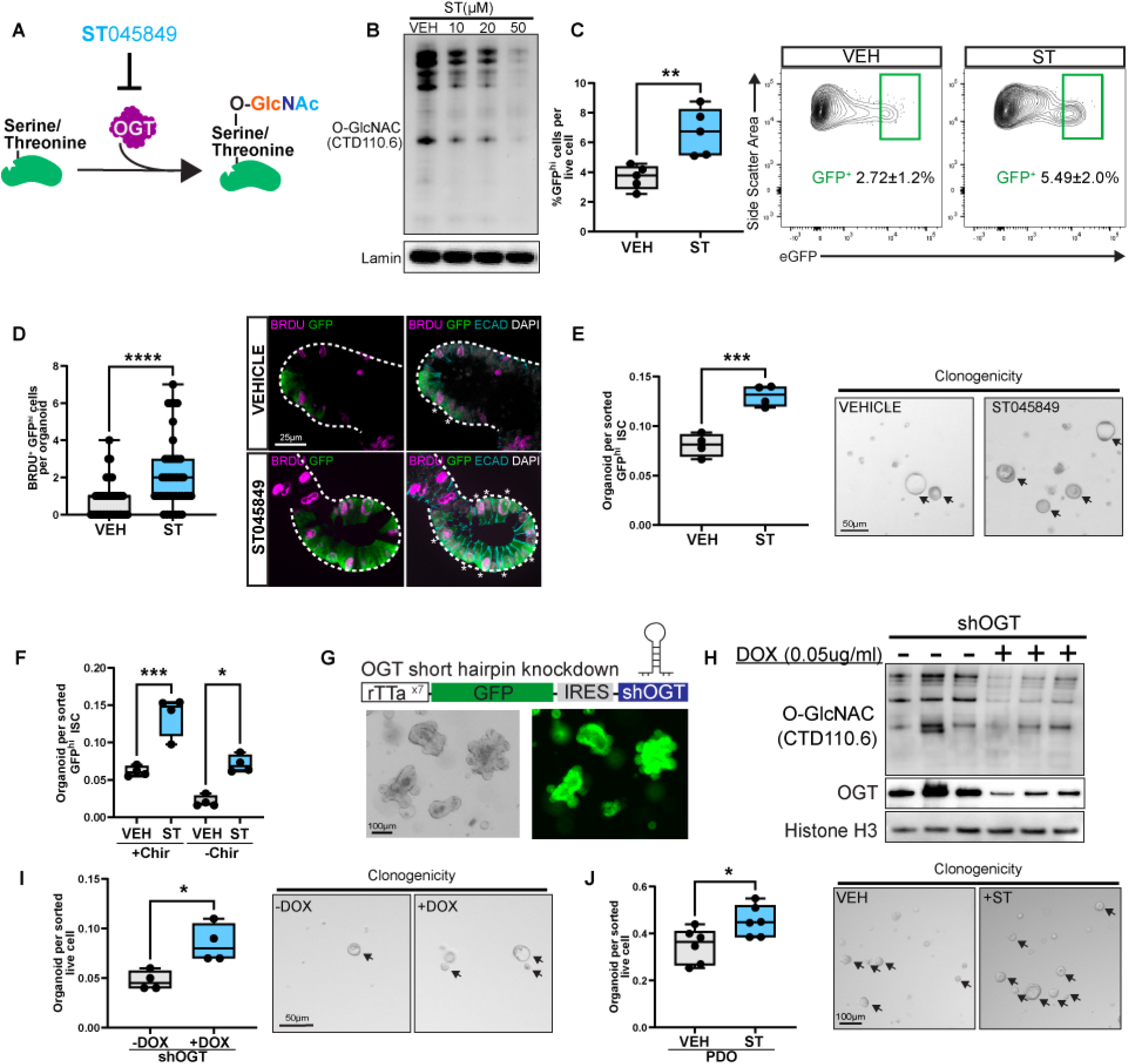
Reduced OGN increases stemness. (A) Schematic of OGT inhibition with pharmacological inhibitor ST045849. (B) Immunoblot analysis of O-GlcNAc (CTD110.6 clone) and Lamin (loading control) in intestinal organoids derived from *Lgr5*^eGFP-IRES-CreERT2^; *Vil*^CreER^ mice treated for 3 days with increasing concentrations of ST. (C) Quantification of GFP^hi^ ISCs in vehicle and ST (20µM) treated organoids and representative flow cytometry plots. N=5, *\*\*p<0*.*01* by Students t-test. (D) Quantification of BrdU^+^GFP^hi^ ISCs per organoid in vehicle and ST treated intestinal organoids and representative immunofluorescent images. Magenta: BRDU; Green: GFP; Cyan: E-cadherin; White: DAPI. N=3, 15 organoids per replicate. *\*\*\*\*p<0*.*0001* by Students t-test. (E) Quantification of clonogenic potential of vehicle and ST treated GFP^hi^ ISCs (flow-isolated) derived from *Lgr5*^eGFP-IRES-CreERT2^; *Vil*^CreER^ mice with representative brightfield images of organoid formation 3 days after plating. N=4. *\*\*\*p<0*.*001* by Students t-test. (F) Clonogenic potential in presence and absence of GSK3b inhibitor, Chir-99021. N=4. \**p<0*.*05* and *\*\*\*p<0*.*001* by two-way ANOVA. (G) Schematic of shOGT intestinal organoid model and representative images of GFP expression post doxycycline administration. (H) Immunoblot analysis of OGT and O-GlcNAc (CTD110.6) in shOGT intestinal oragnoids. (I) Quantification and representative images of doxycycline induced and untreated shOGT organoids N=4. Arrows highlight organoids. \**p<0*.*05* by Students t-test. (J) Quantification and representative images of human jejunum patient derived organoids either vehicle or ST treated N=6. Arrows highlight organoids. \**p<0*.*05* by Students t-test.

To further examine the functional consequences of partial OGT inhibition, we isolated *Lgr5*-eGFP^+^ ISCs from naïve mice, embedded them into Matrigel culture with or without ST, and enumerated the formation of organoids as a measure of self-renewal capacity in a clonogenic assay (Figure 2E). In parallel, we cultured organoids in ST, subsequently collected live epithelial (EPCAM^+^) cells, and similarly tested their clonogenic potential after ST exposure (Figure S2C). Reducing OGN levels enhanced organoid formation in both scenarios, consistent with the expansion of GFP^+^ ISCs and increased proliferation. Given that OGT inhibition confers increased stem cell function, we asked whether ST exposure could compensate for Wnt/β-catenin activity, which is required for ISC maintenance. We found that removing the Wnt activity in isolated *Lgr5*-eGFP^+^ ISC cultures, by excluding the GSK-3b inhibitor (Chir99021), reduced organoid formation (Figure 2F). However, partial OGT inhibition with ST-treatment rescued the growth deficits in absence of this Wnt activity, suggesting that reduced OGN can partially substitute for canonical Wnt input to support ISC self-renewal.

ISCs are known to remodel intestinal composition in response to dietary cues (Beyaz et al., 2016; C. Li et al., 2023; Yilmaz et al., 2012). We tested whether OGN reduction and heightened stemness altered lineage specification. Decreasing OGN increased the abundance of MUC2^+^ goblet cells per organoid, yet, we observed no impact on Paneth (LYZ^+^), tuft (DCLK1^+^), or enteroendocrine (CHGA^+^) cells as assessed by IF. Programmed cell death based on cleaved-caspase3 abundance was also unchanged with reduced OGN (Figure S2D). In addition, we analyzed published scRNA-seq data of jejunum tissue at homeostasis, revealing ISCs and goblet cells have the highest expression of *Ogt*, highlighting the potential for inhibiting OGT to uniquely impact these cell types(Figure S2E) (Enriquez et al., 2022). Overall, reduced OGN selectively biases differentiation toward the goblet lineage in organoids without broadly disrupting epithelial lineage balance or viability.

To confirm that the ISC features were due to partial OGT inhibition and not specific to the ST045849 small molecule, we tested another common OGT pharmacological inhibitor, OSMI-1. Additionally, we interrupted the production of OGT via engineering a doxycycline (dox)-inducible short hairpin targeting the 3’UTR of the transcript (shOGT). We found that both naïve *Lgr5*-eGFP^+^ ISCs treated with OSMI-1 and dox-exposed shOGT organoids demonstrated increased organoid-forming capabilities in the clonogenic assay (Figures 2G-I and S2F-H). We next asked whether the functional effects of OGN modulation are conserved in humans. Using organoids derived from three independent jejunum samples, ST-treatment increased *Lgr5*^+^ cells, seen by RNA single-molecule *in situ* hybridization (smFISH), and enhanced clonogenic potential after sorting the live population and performing organoid-forming assays (Figures 2J and S2I). These findings show that reduced OGN activity, via genetic knockdown and pharmacologic inhibitors, can boost intestinal stemness in a conserved manner, suggesting that glucose derived UDP-GlcNAc and the OGN modification play a role as metabolic regulators of ISC maintenance.

### Decreasing OGN increases PPAR and lipid programs

To understand the extent to which reduced OGN alters metabolic programs, we first performed bulk RNA-sequencing on vehicle- and ST-treated organoids. We observed a robust upregulation of genes involved in lipid metabolism (ie. *Hmgcs2, Pdk4, Me1, Plin2*)(Figure 3A). Gene set enrichment analysis using KEGG enriched for PPAR signaling and fatty acid degradation programs (Figure 3B). Consistent with our analysis of previously published data in other tissues, ST-treatment increased PPAR-dependent transcriptional programs, suggesting that reduced OGN induces a conserved metabolic response (Figure S3A)(Kang et al., 2025). To further assess this relationship, we compared transcriptional changes in ST-treated organoids with those induced by PPARd agonist, GW501516, using RNA-seq. Both conditions produced similar upregulation of lipid metabolic programs, including lipid metabolism, fatty acid oxidation, and lipid droplet biosynthesis, whereas glucose metabolism genes shift more variably in ST-treatment than with the PPARd-agonist (Figure 3C). To quantify the degree of similarity between the PPARd agonist and OGN inhibition in intestinal organoids, we investigated the overlap between differentially expressed genes (DEGs) of the two conditions. We identified 148 DEGs common to both conditions, a highly significant overlap (hypergeometric test, *p* ≈ 1.87 × 10^−216^), indicating substantial concordance in their transcriptional responses (Figure S3B). Moreover, KEGG enrichment and module scoring of the overlapping genes showed conserved transcriptional response of PPAR activation between OGN inhibition and PPARd agonism (Figures S3C-D). We confirmed the increased abundance of lipid mediators and PPAR targets using immunoblots, observing more than 2-fold increases in PDK4, HMGCS2, FABP1, and CPT1a following ST or PPARd agonist treatment (Figures 3D and S3E). Similar increases were observed with the OGT inhibitor OSMI-1 (Figure S3F). Additionally, inhibition of the HBP upstream of UDP-GlcNAc production with azaserine also increases PPAR target HMGCS2 (Figure S3G). Thus far, reduced OGN shifts the ISC metabolism toward a PPAR-driven lipid metabolic program, positioning OGN as a regulator of the metabolic-transcription state in ISCs.

**Figure 3.**
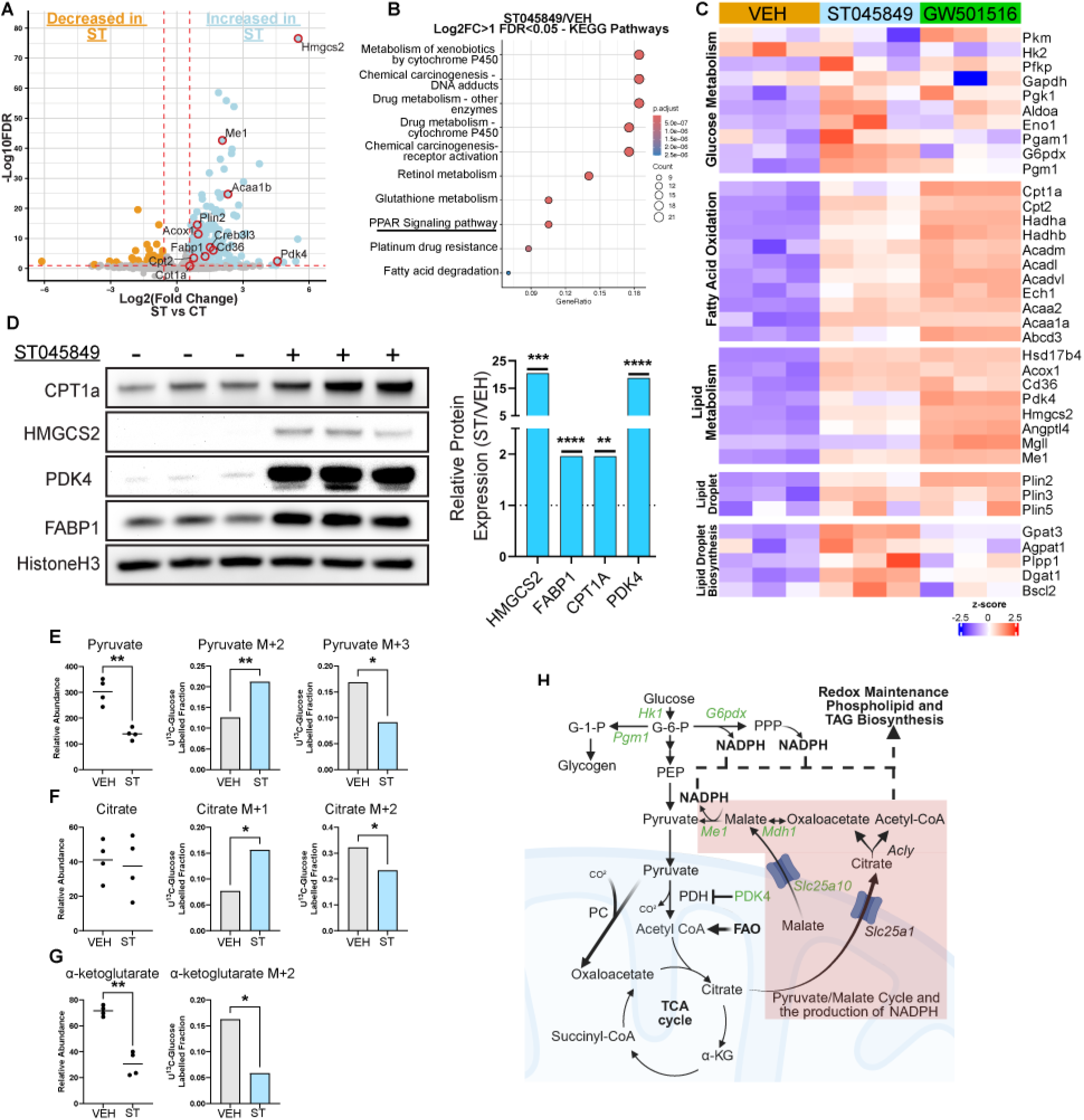
Decreasing OGN increases PPAR and Lipid programs. (A) Volcano plot of comparison between 3day ST and vehicle treated intestinal organoids. Blue: Log2(FC)>0.5 FDR<0.05; Orange: Log2(FC)<-0.5 FDR<0.05. N=3. (B) KEGG enrichment of DEGs in ST treated organoids Log2FC>1 and FDR<0.05. (C) Heatmap of chosen metabolic pathways between vehicle, ST045849 and GW501516 treated intestinal organoids. (D) Immunoblot analysis of PPAR target proteins in vehicle and ST treated intestinal organoids. Quantification of relative protein abundance normalized to HistoneH3 and shown relative to vehicle. N=5 *\*\*p<0*.*01, ***p<0*.*001, ****p<0*.*0001* by Students t-test. (E,F,G) Metabolomic analysis of 24 hour ^U13C^glucose-treated intestinal organoids of relative abundance and mass isotopologue distribution of Pyruvate (E), Citrate (F), and α-ketoglutarate (G). N=4, *\*p<0*.*05 and **p<0*.*01* by Students t-test. (H) Schematic of metabolic changes induced by decreasing OGN in intestinal organoids.

Next, we sought to investigate metabolic rewiring within ISCs and epithelia in reduced OGN conditions. We observed progressive organoid budding morphology over 9 days under partial OGT inhibition, reminiscent of the phenotype in organoids lacking mitochondrial pyruvate carrier, suggesting changes in glucose utilization(Figure S3H) (Schell et al., 2017). Using carbon isotope tracing of ^13^C_6_-glucose, we observe an overall decrease in the abundance of pyruvate in ST-treated organoids, suggesting that decreased OGN moves carbons away from glycolysis and potentially shunts glucose intermediates to other pathways such as pentose phosphate pathway and glycogen, supported by increased transcriptional abundance of *G6pdx* and *Pgm1*, respectively (Figures 3C and 3E). Furthermore, within the pyruvate labeled fractions, pyruvate (M+3) decreased along with TCA intermediates citrate (M+2) and α-ketoglutarate (M+2) (Figures 3E-G). Concomitantly, PDK4, whose activity inhibits 3-carbon pyruvate entering the TCA cycle, also increased in protein abundance, indicating that less OGN allows less pyruvate into mitochondria for oxidation (Figure 3D). Unexpectedly, however, we observed an increase in fractional labeling of pyruvate (M+2)(Figure 3E) coupled with transcriptional abundance of malic enzyme 1 (*Me1*), malate dehydrogenase (*Mdh1*) and mitochondrial dicarboxylate carrier (*Slc25a10*), suggesting that ST-treatment promotes the pyruvate-malate cycle which generates NADPH that can be used for redox balance and lipogenesis (Figures 3C and 3H). To test if diminished OGN increases lipogenesis, we examined lipid droplet accumulation. First, we observed an increase in BODIPY labeling and BODIPY^+^ lipid droplet size in ST-treated MC38 cells (Figure S3I). Second, we saw an increase in PLIN2, a lipid droplet membrane protein and PPAR target, in ST-treated organoids seen by IF and RNA-seq. Both results are consistent with increased lipogenesis and suggest that the pyruvate-malate cycle is a possible metabolic route supporting this shift (Figures 3C and S3J). As a final observation of glucose utilization, we detected increased labeling of citrate (M+1) in ST-treated organoids, which we propose is due to the recycling of ^13^CO_2_ from the increased flux of pyruvate-to-oxaloacetate and malate-to-pyruvate in ST-treated conditions (Figure 3F) (Duan et al., 2022). These data further illustrate that reduced OGN establishes a metabolic state that alters the glucose and lipid oxidation programs to influence ISC function.

### Enhanced ISC function following decreased OGN requires PPARd/a

We next examined whether the observed ST-induced stemness depends on PPAR function. Our previous work showed that simultaneous loss of two nuclear receptors, PPARd and PPARa, blocked the elevated lipid metabolism and stemness induced by a HFD (Beyaz et al., 2016; Mana et al., 2021; Saiz et al., 2026). To test whether these receptors are required for the response to reduced OGN, we derived organoids from intestinal double knockouts of both *Ppars* (*Ppar-d/a*^iKO^: *Ppard*^fl/fl^; *Ppara*^fl/fl^; *Vil*^CreER^; *Lgr5*^eGFP-IRES-CreER^) and wild type controls (WT: *Vil*^CreER^; *Lgr5*^eGFP-IRES-CreER^). We administered ST045849 in culture and examined *Lgr5-*eGFP^hi^ ISCs by flow cytometry, showing that *Ppar-d/a*^iKO^ blocked the increased ISC frequency relative to WT counterparts (Figure 4A). Proliferation, measured through BrdU incorporation, in the ST-treated organoids was also diminished (Figure 4B). Molecularly, we see that *Ppar-d/a* deficiency prevents increased protein levels of PPAR targets HMGCS2 and CPT1a (Figures 4C and S4A). Functionally, *Lgr5-*eGFP^hi^ ISCs derived from *Ppar-d/a*^iKO^ mice failed to exhibit increased regenerative potential as seen in the WT ST-treated controls (Figure 4D). Similarly, clonogenic analysis of WT and *Ppar-d/a*^iKO^ organoids treated for 12 days, thus incorporating 3 passages, showed that the increase in clonogenic potential bestowed by reduced OGN in WT ISCs was abrogated in the *Ppar-d/a*^iKO^ ISCs (Figure S4B). Earlier transcriptional analyses (Figures 3B and S3C) indicated enrichment of fatty acid oxidation programs following partial OGT inhibition, prompting us to test whether the downstream PPAR target, *Cpt1a*, was required for this phenotype. Prior work demonstrates that mitochondrial FAO is required for HFD-driven ISC enhancement, as loss of *Cpt1a* abolishes this phenotype (Mana et al., 2021). Crypts derived from *Cpt1a* conditional knockout mice (*Cpt1a*^iKO^: *Cpt1a*^fl/fl^; *Vil*^CreER^; *Lgr5*^eGFP-IRES-CreER^) and treated with ST over 3 passages similarly failed to show the enhanced clonogenic potential endowed by decreasing OGN (Figures S4C and S4D), highlighting the shift to a PPAR/CPT1a axis with partial OGN inhibition. Furthermore, analysis of published datasets (Mana et al., 2021) shows that loss of *Cpt1a* does not alter *Ogt* expression levels (Figure S4G). In contrast *Ppar-d/a* deficiency increases *Ogt* expression, whereas *Ogt* levels decline under HFD conditions (Figure S4F) implicating a coordinated regulation between OGT and PPAR signaling. As previously established, *Ppard* does not transcriptionally increase in response to conditions that elevate PPAR demand, such as HFD (Beyaz et al., 2016) or fasting (Figures S4H and S4I) (Mihaylova et al., 2018), suggesting that enhanced activity is regulated primarily through cofactors and post-translational modifications rather than transcriptional control.

**Figure 4.**
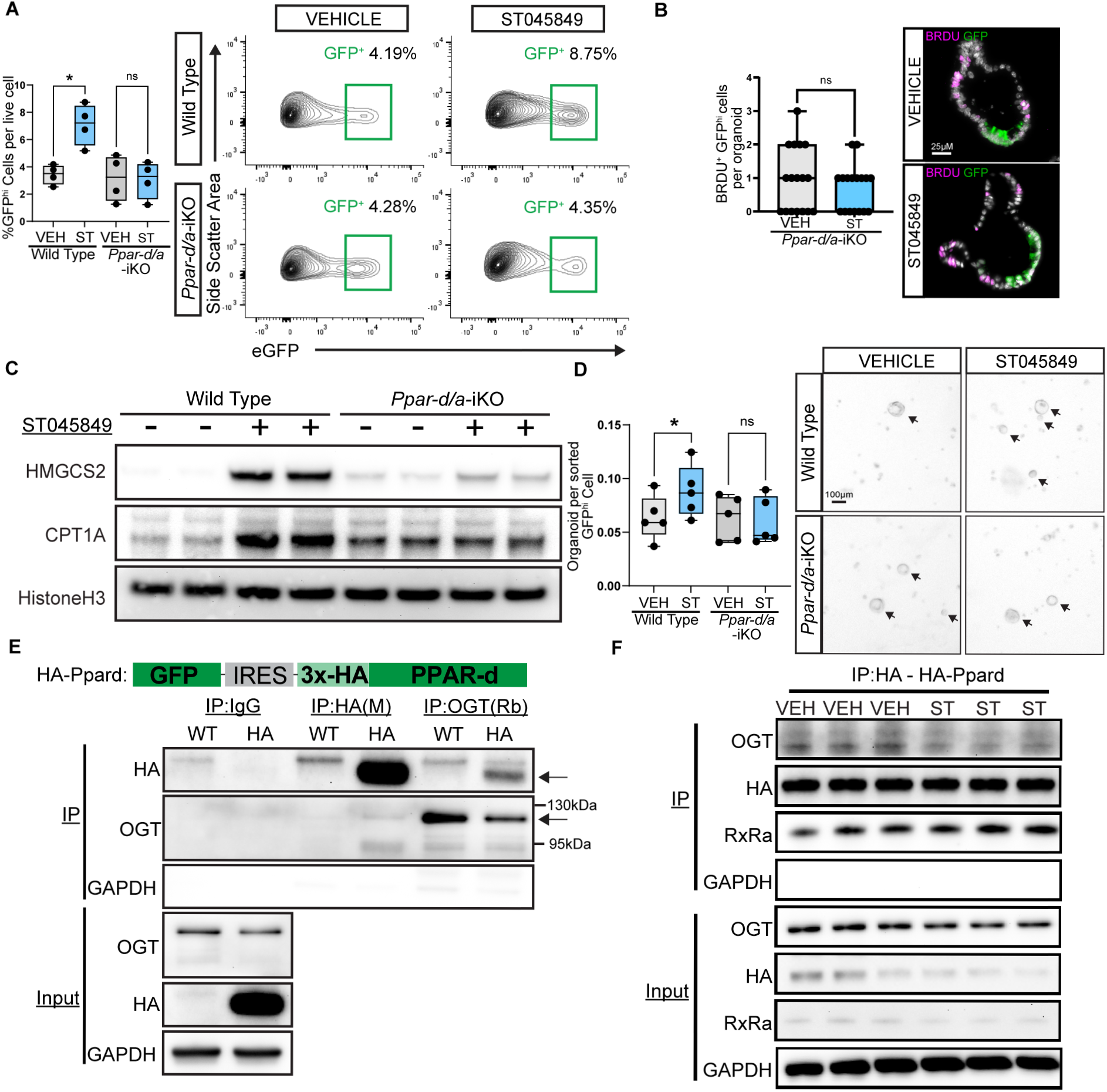
Enhanced ISC function following decreased OGN requires PPAR*d/a*. (A) Quantification and representative flow plots of *Lgr5*-eGFP^hi^ ISCs of vehicle and ST-treated organoids derived from (*Ppar-d/a*^iKO^: *Ppard*^fl/fl^; *Ppara*^fl/fl^; *Vil*^CreER^; *Lgr5*^eGFP-IRES-CreER^) and wild type controls (WT: *Vil*^CreER^; *Lgr5*^eGFP-IRES-CreER^). N=4, ns: not significant, *\*p<0*.*05* by two-way ANOVA. (B) Quantification of BRDU^+^GFP^hi^ ISCs per organoid in vehicle and ST treated *Ppar-d/a*^iKO^ intestinal organoids and representative immunofluorescent images Magenta: BRDU. Green: GFP, White: DAPI. N=3. ns: not significant by Students t-test (C) Immunoblot of PPAR target proteins in WT and *Ppar-d/a*^iKO^ intestinal organoids treated with ST. (D) Quantification and representative images of organoids formed per sorted *Lgr5*-eGFP^hi^ ISC from WT and *Ppar-d/a*^iKO^ animals treated with either vehicle or ST. N=5. ns: not significant, *\*p<0*.*05* by two-way ANOVA. (E) Immunoblot analysis of input and anti-HA, anti-IgG and anti-OGT immunoprecipitate from WT and HA-PPARd expressing MC38 cells. (F) Immunoblot analysis of input and anti-HA immunoprecipitate of HA-PPARd MC38 cells treated with vehicle or ST.

### PPARd and OGT form a nexus of lipid and sugar metabolic regulation

The data highlighted that the PPAR/CPT1a axis rapidly responds to partial OGT inhibition, leading us to question whether OGT directly modulates PPARd and that the lack of this modification stimulates PPAR activation and subsequent lipid metabolism. In absence of reliable antibodies against PPARd, we generated a 3xHA-tagged *Ppar-d* expression construct (HA-PPARd) and subsequently transduced it into a genetically *Ppard*-deficient MC38 cell line (*Ppard-*KO) to allow the immunoprecipitation of tagged HA-PPARd from near endogenous levels. We tested these cells to confirm PPAR target proteins remain similar to WT and that ST treatment could elevate target protein abundance (HMGCS2, CPT1a, ECH1)(Figure S4E). Additionally, we confirmed that immunoprecipitation of PPARd pulls down its obligate binding partner RXRa (Figure 4F). We hypothesized that if OGT directly modifies PPARd, we should be able to detect OGT in an HA-PPARd immunoprecipitant. Indeed, we found that OGT binds to HA-PPARd, and conversely, IP of OGT pulls down HA-PPARd confirming these two proteins can complex together (Figures 4E and Figure S4J). Moreover, in ST-treated cells, we observed decreased OGT levels in HA-PPAR IPs, substantiating a role of OGT in binding PPARd (Figure 4F). Together, these data reveal a dynamic OGT–PPARd interaction that may couple nutrient-sensitive O-GlcNAc signaling to PPAR-driven transcriptional programs.

## DISCUSSION

Here we propose that ISCs dynamically coordinate fuel utilization through an OGN–PPAR regulatory axis that facilitates transitions between glucose- and lipid-based metabolism. We demonstrate a direct interaction between OGT and PPARd, and observe reduced complex formation under partial OGT inhibition coincident with enhanced PPAR activity, altered pyruvate oxidation, and augmented stemness. Our findings support a model in which OGN-PPAR functions as a tunable molecular rheostat that modulates lipid transcriptional programs in response to the nutrient state.

Under homeostatic conditions, glucose is likely a primary fuel source for ISCs (C. Li et al., 2023; Ludikhuize et al., 2020; McCauley et al., 2023; Rath et al., 2018b; Rodríguez-Colman et al., 2017). In this context, we suggest that OGN functions as a glucose-responsive brake on PPAR activity. Because OGN depends on UDP-GlcNAc derived from glucose metabolism, high-glucose states may promote OGN of PPAR or its associated cofactors, thereby maintaining PPAR activity at a relatively subdued level. This would favor glucose utilization and constrain lipid-driven transcriptional programs. In contrast, when OGT activity is reduced, either pharmacologically, through dietary shifts that alter metabolite availability, or upon differentiation with more demands for PPAR activity, OGN levels decline thereby releasing this restraint and permitting enhanced PPAR activation. This shift promotes lipid metabolic programs and mitochondrial fatty acid oxidation in ISCs. Rather than viewing glucose and lipid metabolism as independent pathways, our work suggests that ISCs respond to dietary metabolic availability by integrating glucose flux with lipid transcriptional control along an OGN-PPAR axis, enabling ISCs to dynamically shift between fuel sources. Furthermore, this mechanism resolves longstanding observations of altered PPARd activity without changes in *Ppard* transcriptional abundance, suggesting protein-level regulation as a dominant control point.

We distinguish between partial inhibition of OGT and complete loss of OGT function. Because OGT is the sole enzyme responsible for catalyzing OGN, complete inhibition or genetic loss of OGT would be expected to produce widespread metabolic disruption (X. Li et al., 2023; Shafi et al., 2000; Yang et al., 2020; Zhao et al., 2018). In contrast, the consistent and robust phenotype observed with partial OGT inhibition suggests a specific regulatory role rather than a nonspecific metabolic consequence. Complete intestinal deficiency of OGT *in vivo* results in an expansion of the crypt domain and increases the number of proliferating cells; however, this is accompanied by increased apoptosis, reduced barrier integrity, and elevated inflammation (Zhao et al., 2022, 2020, 2018). Crypt-derived OGT-knockout organoids fail to thrive (Zhao et al., 2020, 2018). These phenotypes are inconsistent with high PPAR activity and instead reflect the broader cellular dysfunction that arises from complete OGN loss. Our work suggests that one major outcome of OGN is the fluctuating control of fuel utilization and may represent a broader paradigm for how metabolic signals are translated into transcriptional programs.

## EXPERIMENTAL MODEL AND STUDY PARTICIPANT DETAILS

Mice were under the husbandry care administered by the Department of Animal Care and Technologies at Arizona State University (ASU). All procedures were conducted in accordance with the American Association for Accreditation of Laboratory Animal Care and approved by ASU’s Institutional Animal Care and Use Committee. The following alleles were obtained from the Jackson Laboratory or gifted: *Rosa26*^LSL-3XHA-EGFP-OMP25^ (B6.Cg-Gt(ROSA)26Sor^tm1(CAG-EGFP)Brsy^*/*J, Jax # 032290)(Bayraktar et al., 2019);

*Lgr5*^CreERT2^ (Huch et al., 2013); *Rosa26*^LSL-tdTomato-fl/fl^ (B6.Cg-Gt(ROSA)26Sor^tm9(CAG-tdTomato)Hze^/J,Jax # 007909); *Ppard*^fl/fl^ (B6.129S4-Ppard^tm1Rev^/J, Jax # 005897)(Barak et al., 2002); *Ppara*^fl/fl^ (Brocker et al., 2017); *Lgr5*^eGFP-IRES-CreERT2^ (B6.129P2-Lgr5^tm1(cre/ERT2)Cle^/J)(Barker et al., 2007); *Cpt1a*^fl/fl^ (Schoors et al., 2015); *Villin*^CreERT2^ (B6.Cg-Tg(Vil1-Cre/ERT2)23Syr/J, Jax # 020282)(el Marjou et al., 2004). The following strains were bred in-house: (1) WT: *Villin*^CreER^; *Lgr5*^eGFP-IRES-CreERT2^, (2) *Ppar-d/a*^iKO^:*Ppard*^fl/fl^; *Ppara*^fl/fl^; *Villin*^CreER^; *Lgr5*^eGFP-IRES-CreERT2^, (3) *Cpt1a*^iKO^: *Cpt1a*^fl/fl^; *Villin*^CreER^; *Lgr5*^eGFP-IRES-CreERT2^, (4) MITO-Tag^ISC^: *Lgr5*^CreERT2^; *Rosa26*^LSL-3XHA-EGFP-OMP25^, (5) *Lgr5*^CreERT2^; *Rosa26*^LSL-tdTomato-fl/fl^. Mice were sex and age-matched per experiment and housed under a 12/12 day/night light cycle. Floxed alleles were excised following intraperitoneal injection of tamoxifen administered at 50-100 mg/kg. Mice harboring *Cpt1a* or *Ppar* alleles were administered tamoxifen 3 times at 100, 50 and 50 mg/kg over the course of two weeks prior to use for harvesting organoids. High-fat diet (Research Diets D12492) and Control diet (matching sucrose, Research Diets D12450J) were provided to mice *ad libitum* at the age of 8-12 weeks for 1month. Fasted mice were moved to a clean cage with access to water and no access to food at 9 AM. Mice were sacrificed 24 hours later.

## METHOD DETAILS

### Crypt Isolation and establishment

Small intestines were removed, flushed with PBS^-/-^ (No Ca^2+^, No Mg^2+^), opened laterally and cut into 3-4cm pieces after removal of the mucus layer. Intestines were rinsed with 25mL of PBS^-/-^ and incubated for 35 min on a rocker at 4°C in PBS^-/-^ + 10mM EDTA (Invitrogen^TM^, 15575020). By mechanical shaking crypts, were removed from connective tissue and filtered through 70μm mesh to isolate crypts from villi and tissue fragments. Isolated crypts were then pelleted at (300 g, 5 min, 4°C) and used for isolation of ISCs (described below) or counted and resuspended in a 35:65 ratio of media: Matrigel (Corning 356231 growth factor reduced) at 12 crypts per μL and plated in 10μL domes. After a 15minute incubation at 37°C, media was added. Unless otherwise described crypts were grown in RPMI-1640 (Gibco^TM^, 11879020) supplemented with Glucose 5mM, 1x Penicillin-Streptomycin (Gibco^TM^,15140149), mEGF 40ng/mL (Peprotech, 315-09), 1x B27 (Life Technologies, 17504044), N-acetyl-L-cysteine 1 μM (Sigma Aldrich, A9165), Chir99021 1 μM (LC Laboratories, C-6556), Y-27632 dihydrochloride monohydrate 10 μM (Sigma Aldrich, Y0503) and conditioned media corresponding to 200ng/ml Noggin (HEK293-mNoggin-Fc)and 500ng/mL R-spondin (HA-R-Spondin1-Fc, R&D systems, 3710-001-01). Intestinal organoids were maintained at 37°C in a humidified incubator at 5% CO_2_. Media was changed every 3 days as appropriate. ST045849 (Sigma Aldrich, SML2702) and GW501516 (Sigma Aldrich, SML1491) were added at 20μM and 1μM, respectively, unless otherwise stated. For U-^13^C_6_-Glucose tracing the same media as above was used except with the replacement of media 24hours prior to collection with D-Glucose (U-^13^C_6_, 99%, Cambridge Isotope Laboratories, Inc., CLM-1396). For BrdU incorporation 20 μM was administered to organoids for 20 min at 37°C prior to collections.

### Isolation of ISCs and Flow Cytometry

Following crypt isolation, crypts were resuspended in TrypLE (Gibco^TM^, no phenol red, 12604039) and dissociated into single cells through a 60-second incubation at 32°C. Dissociated single cells were washed with 10ml of S-MEM (Gibco^TM^, 11380037) and pelleted (300 g, 5 min, 4°C). Following resuspension and filtering through a 70μm mesh cells were incubated with EpCAM-APC (Bioscience, 17-5791-81) and DAPI (BioLegend, 422801) was used to exclude dead cells from analysis. ISCs were isolated as Lgr5-eGFP^hi^EpCAM^+^ cells and TA progenitor cells were isolated through Lgr5-eGFP^low^EpCAM^+^ using BD FACS Symphony S6. Isolated Lgr5-eGFP+ cells were centrifuged for 5minutes at 300g, resuspended in an appropriate volume of crypt medium, and seeded onto 5 μL Matrigel (Corning 356231 growth factor reduced) containing 1 μM JAG-1 (Anaspc, AS61298) in a flat bottom 96-well plate (Olympus, 25-109). Clonogenic capacity was assessed after 3-days, and brightfield images were captured using a Keyence BZ-X810.

For flow cytometry isolation of single cells from organoids, 3 days after initial crypt plating, organoids were collected in PBS^-/-^ and washed to remove Matrigel matrix. Organoids were pelleted through 3-4 rapid 3-second centrifugation steps and the supernatant was removed. The organoid pellet was resuspended in 300 μL of TrypLE and incubated for 3minutes at 37°C to dissociate into single cells. Single cells were then washed and incubated with EpCAM-APC (Bioscience, 17-5791-81) and DAPI (BioLegend, 422801) was used to exclude dead cells from analysis. Clonogenic capacity and plating of single cells were completed as above for Lgr5+ ISCs of LIVE^+^EpCAM^+^ cells.

### Embedding and Immunofluorescence (IF)

Tissues were fixed in 10% formalin, paraffin-embedded and sectioned in 4-5micron sections as previously described (Yilmaz et al., 2012). Embedding and sectioning were done by the Mayo Clinic Research Histology Core. Organoids were collected and fixed in 4% PFA (Pierce^TM^, 28906) for 15minutes followed by washing in PBS^-/-^. Organoids were resuspended in HistoGel™ (Epredia™, 22-110-678) and paraffin-embedded and sectioned in 4-5micron sections. Antigen retrieval was performed using Borg Decloaker RTU solution (Biocare Medical, BD1000G1) and a pressurized Decloaking Chamber (Biocare Medical, NxGen). Antibodies and respective dilutions used for immunofluorescence are as follows: Mouse monoclonal anti-CPT1A (1:250, abcam 128568), Goat polyclonal anti-GFP (1:500, abcam 6673), Rabbit monoclonal anti-GFP (1:500, CST 2956), Rabbit monoclonal anti-OLFM4 (1:1000, CST 39141), Rabbit monoclonal anti-Cleaved Caspase 3 (1:1000, CST 9664), Rat monoclonal anti-BrdU (1:500, abcam 6326), Rabbit polyclonal anti-Lysozyme (1:500, ThermoFisher RB-372-A1), Rabbit polyclonal anti-ChromograninA (1:500, Abcam 15160), Rabbit monoclonal anti-DCAMLK1 (1:500, Abcam 109029), Rabbit monoclonal anti-HA-Tag (1:500, CST 3724), Rabbit polyclonal anti-MUC2 (1:500, Novus Biologicals NBP1-31231). Alexa Fluor secondary antibodies, anti-mouse 488, anti-rabbit 568, anti-rat 647, and anti-goat 647 were used at 1:500. Tissue and organoids were mounted in Invitrogen Prolong Gold Antifade with DAPI (Invitrogen, P36931). Images were acquired on a Keyence BZ-X800. For imaging of mitochondria and HA-tag in Supplemental Figure 1 images were acquired on a Zeiss 880 confocal with Airyscan.

For *ex vivo* imaging of whole-mount tdTomato tissue 40 hours post tamoxifen induction, small intestine tissue was extracted, washed, and opened laterally. Tissue was then pinned on 35mm dishes and imaged in RPMI 1640 media with using a STELLARIS DIVE Multiphoton Microscope with dipping objective lens.

### Single-molecule fluorescent in situ hybridization (smFISH)

smFISH was performed using Advanced Cell Diagnostics RNAScope™ Multiplex Fluorescent Reagent Kit v2 and according to the manufacturer’s instructions. The smFISH probes used was the Hs-LGR5-C2 (Ref 311021-C2). Analysis was conducted blind and done using Qupath. >10 organoids were analyzed per replicate. Cells with >5 puncta were considered Lgr5^+^ cells.

### Patient-Derived Organoid Culture

Human jejunal organoids were cultured using previously defined media and methods with minor modifications (Miyoshi and Stappenbeck, 2013). Human jejunal media composed of: 50% L-WRN conditioned media and Advanced DMEM/F12 (Gibco, 12634010) supplemented with 1% GlutaMAX-I (Gibco, 35050-061), 1% PenStrep (Gibco, 15140-122), 10 mM HEPES, 50 ng/mL EGF (Life Technologies, PHG0311), 1 mM N-acetyl-l-cysteine (Sigma, A9165), 1x N2 (Life Technologies, 17502-048), 1x B27 (Life Technologies, 12587-010), 10 nM [Leu15]-Gastrin 1 (Sigma, G9020), 10 mM Nicotinamide (Sigma, N3376), 500 nM A83-01 (Tocris, 2939), 3 μM SB202190 (Sigma, S7067), 100 μg/mL primocin (Invivogen, #ant-pm), 10 μM Y-27632 (Sigma, Y0503). For differentiation 50ng/ml of FGF-2 (R&D Systems 3718-FB) and 100ng/ml of IGF-1 (R&D Systems, 291-G1) was added to the media. Organoid media was changed every 3-days. Post passaging, organoids were plated with differentiation media and treated for 3-days prior to collection with ST045849.

### Electron Microscopy

Tissue was collected and chemically fixed for 24 hours at room temperature with 2.5% glutaraldehyde in cacodylate buffer. Samples were then washed 3 times with cacodylate buffer followed by secondary fixation in 1% osmium in 0.1M cacodylate and 2% aqueous uranyl acetate. Tissue was embedded in Spurr’s resin, and ultra-thin sections were cut and transferred onto Formvar-coated copper grids. STEM images were collected on the Phenom Pharos Desktop STEM.

### MITO-Tag Mitochondrial Isolation

MITO-Tag mitochondrial immunopurification was performed as previously described, with certain modifications (Bayraktar et al., 2019). All steps for mitochondrial isolation were performed on ice using pre-chilled buffers. Intestinal tissue was extracted, flushed with KPBS (136 mM KCl, 10 mM KH2PO4, pH 7.25), opened laterally, and villi were removed through scraping with a glass slide. Tissue (≈100mg) was minced prior to being transferred to a Dounce homogenizer with 1.5mL of KPBS containing protease and phosphatase inhibitors (Thermo Scientific^TM^, 78441), and homogenized for 25 strokes. 25 μL of the whole-cell lysate was collected for downstream analysis. The remaining homogenate was spun down at 1,000 x *g* for 2 minutes at 4°C. The supernatant was then collected, 25 μL was collected again as the input for downstream analysis, and the remaining supernatant was incubated with anti-HA magnetic beads (Pierce^TM^, 88836) pre-washed with KPBS on a spinning rotator for 10minutes at 4°C. 3 rounds of washing with KPBS was followed by 10% of the suspension being separated and used for protein extraction and immunoblotting. The remaining suspension was collected in in ice cold 80:20 MeOH and LC/MS Grade H_2_O (Millipore Sigma, 1153331000) for downstream analysis.

### Cell Lines and Generation of Genetically Edited Organoids

The shOGT construct was based on LT3GECIR (Johannes Zuber: Addgene #111178), with shRNA designed using splashRNA and inserted to make pLenti-TetON GFP-shOGT puro rTta (Ben Bartelle: Addgene #254223). Lentiviral based gene delivery into C57B6J derived organoids was performed as previously described protocol (Maru et al., 2016).

The lentiviral vector with 3XHA tagged PPARd (HA-PPARd) was designed with the pLenti6 vector with all elements synthesized and assembled into CMV GFP IRES 3XHA-PPARd Puro (Ben Bartelle: Addgene #254211). MC38-*Ppard-*KO and parental wild-type MC38 cell lines were obtained from Ubigene. Cells were maintained in 90% DMEM (Gibco^TM^, 11995065), 10% FBS (Gibco^TM^, A5670801), with 1x Penicillin-Streptomycin (Gibco^TM^,15140149).

### Immunoprecipitation

Immunoprecipitation was performed following the manufacturer’s instructions of the Pierce™ Classic Magnetic IP/Co-IP Kit (Thermo Scientific^TM^, 88804) with the following modifications: PugNAc and Thiamet-G at 10 μM were added to the lysis buffer and wash steps to inhibit OGA activity. Antibodies used for immunoprecipitation were Rabbit monoclonal anti-HA-Tag (1:50, CST 3724) and rabbit polyclonal anti-OGT (1:100, ProteinTech 11575-2-AP). Cell lysate was incubated overnight at 4°C with the antibody followed by 1 hour Room temperature incubation with pre-washed magnetic beads. Beads were washed 5 times with IP Lysis/Wash Buffer containing inhibitors as above, followed by a final wash with ultra-pure water. Samples were eluted from beads using 1X Laemmli buffer (Thermo Scientific Chemicals, J61337.AD) at 95°C for 10 minutes.

### Immunoblotting

Either 10,000 sorted cells were loaded per sample or 10 µg of protein for cell line lysates and tissue lysates were loaded on a 4-12% gradient gel (Invitrogen^TM^, NP0335) and transferred on to PVDF membrane (Immobilon-P transfer, Millipore, ipvh00010).The following antibodies were used: mouse monoclonal anti-Cpt1a (1:500(sorted), (1:1000 (lysates), Abcam ab128568), rabbit monoclonal anti-HMGCS2 (1:500(sorted), Abcam ab137043), rabbit monoclonal anti-HA-Tag (1:1000, CST 3724) and rabbit polyclonal anti-OGT (1:500, Protein Tech 11575-2-AP), rabbit monoclonal anti-FABP1 (1:1000, CST, 13368), monoclonal rabbit anti-PDK4 (1:1000, Abcam, ab214938), rabbit monoclonal anti-total H3 (1:3000, CST 4449S), rabbit monoclonal anti-LAMP1 (1:1000, abcam 208943), rabbit polyclonal anti-Catalase (1:1000, abcam 16731), rabbit monoclonal anti-calreticulin (1:1000, CST 12238), rabbit monoclonal anti-GAPDH (1:1000, CST 2118), mouse monoclonal anti-COXIV (1:1000, CST 11967), rabbit monoclonal anti-TOM20 (1:1000, CST 42406), rabbit monoclonal anti-LAMINB1 (1:2000, Abcam 133741), mouse monoclonal anti-O-GlcNAc (CTD110.6, 1:1000, CST 9875) and mouse monoclonal anti-O-GlcNAc (RL2, 1:1000, ThermoFisher MA1-072). Samples were visualized using IgG-HRP antibodies (1:3000, CST, 7076, 7074) and SuperSignal™ West Pico or Femto (Thermo Scientific^TM^, 34580 and 34096).

### Metabolomics, 13^C^C-Glucose tracing, and LC/MS Methods

Using stable isotopic tracers, metabolic flux analysis was performed according to previously reported methods with minor modifications (He et al., 2020; Shi et al., 2020; He et al., 2023). Briefly, intestinal organoids cultured as described above were switched to fresh RPMI crypt media containing U-^13^C_6_-Glucose 24 hours prior to collection in ice cold 80:20 MeOH and LC/MS Grade H_2_O (Millipore Sigma, 1153331000).

Fluxes of citrate, α-ketoglutarate, pyruvate and additional TCA cycle and amino acid metabolites were measured using GC-MS. After drying, the samples were incubated with a solution of 40 μL of 20 mg/mL O-methylhydroxylamine hydrochloride in pyridine at 60 ^o^C for 90 min. Next, 70 μL of N-tert-butyldimethylsilyl-N-methyltrifluoroacetamide (MTBSTFA) was added and incubated at 60°C for 30 min. An Agilent 8860 GC-5977 MSD system (Agilent Technologies, Inc., Santa Clara, CA, USA) was used for GC-MS spectral acquisition. In addition, fluxes of glycolytic metabolites were measured using LC-MS following previously established protocols (Gu et al., 2015; He et al., 2020; Shi et al., 2020; Wei et al., 2021; He et al., 2023). All LC-MS experiments were performed on a Thermo Vanquish UPLC-Exploris 240 Orbitrap MS instrument (Waltham, MA). Chromatographic separations were performed in hydrophilic interaction chromatography (HILIC) mode on a Waters XBridge BEH Amide column (150 x 2.1 mm, 2.5 µm particle size, Waters Corporation, Milford, MA). The mobile phase was composed of Solvents A (10 mM ammonium acetate, 10 mM ammonium hydroxide in 95% H_2_O/5% ACN) and B (10 mM ammonium acetate, 10 mM ammonium hydroxide in 95% ACN/5% H_2_O). After the initial 1 min isocratic elution of 90% B, the percentage of Solvent B decreased to 40% at t=11 min. The composition of Solvent B maintained at 40% for 4 min (t=15 min), and then the percentage of B gradually went back to 90%, to prepare for the next injection. Using mass spectrometer equipped with an electrospray ionization (ESI) source, we will collect untargeted data from 70 to 400 m/z. After data collection, the mass isotopomer distributions (MIDs) for each sample were calculated by integrating individual metabolites, and the IsoCor software was employed for the correction of natural abundance of isotopes.

To measure metabolite abundances in MITO-tag samples, the targeted LC-MS/MS method used here was modeled after that developed and used in a growing number of studies (Carroll et al., 2015; Gu et al., 2016; Jasbi et al., 2019; Shi et al., 2019; Eghlimi et al., 2020). Briefly, all LC-MS/MS experiments were performed on an Agilent 1290 UPLC-6490 QQQ-MS (Santa Clara, CA) system. Each sample was injected twice, 10 µL for analysis using negative ionization mode and 4 µL for analysis using positive ionization mode. Both chromatographic separations were performed in hydrophilic interaction chromatography (HILIC) mode on a Waters XBridge BEH Amide column (150 x 2.1 mm, 2.5 µm particle size, Waters Corporation, Milford, MA). The flow rate was 0.3 mL/min, auto-sampler temperature was kept at 4°C, and the column compartment was set at 40°C. The mobile phase was composed of Solvents A (10 mM ammonium acetate, 10 mM ammonium hydroxide in 95% H_2_O/5% ACN) and B (10 mM ammonium acetate, 10 mM ammonium hydroxide in 95% ACN/5% H_2_O). After the initial 1 min isocratic elution of 90% B, the percentage of Solvent B decreased to 40% at t=11 min. The composition of Solvent B maintained at 40% for 4 min (t=15 min), and then the percentage of B gradually went back to 90%, to prepare for the next injection. The mass spectrometer is equipped with an electrospray ionization (ESI) source. Targeted data acquisition was performed in multiple-reaction-monitoring (MRM) mode. The whole LC-MS system was controlled by Agilent Masshunter Workstation software (Santa Clara, CA). The extracted MRM peaks were integrated using Agilent MassHunter Quantitative Data Analysis (Santa Clara, CA).

Mitochondrial metabolites were determined as previously described (Bayraktar et al., 2019). MetaboAnalyst 6.0 was used for processing mitochondrial metabolite peak intensity data. Following log transformation and auto-scaling, the normalized abundance of metabolites was analyzed for significance using one-way ANOVA. The MetaboAnalyst Quantitative Enrichment Analysis module was performed on a concentration table of the Control and HFD mitochondrial metabolites.

### RNA-seq data processing and differential expression analysis

Organoid samples were collected in 400 µL of TRIzol^TM^ Reagent (Invitrogen^TM^, 15596026). RNA libraries were sequenced on an Illumina NovaSeq platform to generate 150-bp paired-end reads (PE150). Raw sequencing reads were assessed for quality using FastQC, and adapter sequences and low-quality bases were trimmed using Trim Galore with default parameters. Cleaned reads were aligned to the mouse reference genome (GRCm39) using HISAT2 (v2.0.5) with default settings and gene annotation from GENCODE. Only uniquely mapped reads were retained for downstream analyses. Gene-level read counts were generated using FeatureCounts (v1.5.0-p3) was used followed by FPKM calculation.

Differential gene expression analysis was performed in R using DESeq2. Genes with low expression (total counts <10 across all samples) were excluded prior to analysis. Count data was normalized and genes with a false discovery rate (FDR) < 0.05 and absolute log2 fold change > 1 were considered significantly differentially expressed genes (DEGs). Volcano plots and heatmaps were generated using normalized expression values to visualize differential expression patterns. Kyoto Encyclopedia of Genes and Genomes (KEGG) pathway enrichment analysis was conducted in R using clusterProfiler based on significantly up- and downregulated DEGs. The statistical significance of overlapping DEGs between two conditions was assessed using a hypergeometric test implemented in R (phyper function), with the background defined as all genes. Module scores based on selected DEG gene sets were calculated by averaging normalized expression values of genes within each set for each sample to represent coordinated gene program activity, and statistical comparisons between groups were performed as indicated in the corresponding figure legends.

## QUANTIFICATION AND STATISTICAL ANALYSIS

Graphpad Prism^TM^ was utilized for statistical analysis. Students t-test, one-way ANOVA and two-way ANOVA were used where appropriate and at least three replicates were used for each experiment.

## ACKNOWLEDGMENTS

Cell sorting was assisted by Adam Kindelin and the ASU Biosciences Flow Cytometry Core using BD FACSymphony S6, acquired by the NIH SIG award 1-S10-OD032287-01. The authors acknowledge support from Dr. Honor Glenn at the ASU Biodesign Advanced Light Microscopy Facility. ASU Research Computing provided HPC and storage support (Jennewein et al., 2023). C.F. acknowledges the Wayne State University Karmanos Cancer Institute’s Cancer Center Support Grant (P30CA022453) and NIH/NIGMS R35GM142637 for the purchase of reagents and proteomics support, as well as NIH/NIEHS R01ES035692 for supporting C.A.’s salary and tuition. B.B.B. is supported by NIBIB-R21EB034970, NIMH-DP2MH136493. M.D.M. is supported by NCI-K22CA241083, RSG-22-095-01-CCB, NCI-R01CA301086, NIDDK-R01DK143944.

## AUTHOR CONTRIBUTIONS

Conceptualization, M.D.M., T.H.M., B.B.B.; methodology, M.D.M., T.H.M., B.B.B.; Investigation, T.H.M., Y.B.M., D.R.S., G.L., E.U., C.A., N.H., F.F., J.C., H.G. and C.F.; writing—original draft, T.H.M.; writing— review & editing, M.D.M., B.B.B., and all authors; funding acquisition, M.D.M.; supervision, M.D.M. and B.B.B.

