## Supplemental Figure Legends for "O-GlcNAcylation regulates PPAR-driven metabolic programming in intestinal stem cells"

**Supplemental figure 1. MITO-Tag reveals decreased UDP-GlcNAc in HFD Mitochondria**

(A) *Rosa26*$^{tdTomato}$; *Lgr5*$^{CreERT2}$ jejunum tissue 40- or 60-hours post 100 mg/kg Tamoxifen administration. Red: Tdtomato/RFP; White: DAPI.

(B) *Ex vivo* live images of whole-mount *Rosa26*$^{tdTomato}$*; Lgr5*$^{CreERT2}$ 40-hours post tamoxifen administration. Red: Tdtomato. White: Collagen through fluorescence lifetime imaging microscopy.

(C) Representative fluorescent images of MITO-Tag$^{ISC}$ mice either not induced with tamoxifen (CONTROL-MITO) or 1 month Control and HFD mice 40-hours post Tamoxifen administration. Magenta: HA-MITO; Green: OLFM4; White: DAPI.

(D) Transmission electron microscopy images of intestinal stem cell and organelle contact sites. Black arrow: mitochondria; Red arrow: endoplasmic reticulum; Orange arrow: peroxisome.

(E) Volcano plot of mitochondrial metabolites of the Log2(FC) of HFD mitochondrial metabolites relative to Control.

(F) Normalized abundance of N-Acetylneuraminic acid, a downstream metabolite from UDP-GlcNAc, from Control, HFD and Fasted MITO-Tag$^{ISC}$ mice. ***p<0.01,* one-way ANOVA.

**Supplemental figure 2. Reduced OGN increases stemness**

(A) Immunoblot analysis of O-GlcNAc (CTD110.6) after 3 days of glucose treatment at 5 mM, 15 mM and 25 mM.

(B) Representative images of *Lgr5*$^{eGFP-IRES-CreERT2}$; *Vil*$^{CreER}$ intestinal organoids 3 days post treatment with increasing concentrations of ST045849.

(C) Quantification of clonogenic potential of sorted live cells from vehicle and ST045849 treated organoids. N=5. **p<0.05* by Student's t-test.

(D) Cell type quantification and representative images of vehicle and ST045849 treated organoids. Yellow stars highlight MUC2$^{+}$ goblet cells. Red: Cell specific marker; White: DAPI. N=3, 15+ organoids per replicate. *****p<0.0001 and* ns: not significant by Student's t-test.

(E) Violin plotting of log-normalized *Ogt* expression across selected cell types from GSE199776. Numeric FDR shown for comparisons between ISCs and other cell types.

(F) Representative images of *Lgr5*$^{eGFP-IRES-CreERT2}$; *Vil*$^{CreER}$ vehicle, ST045849 or OSMI-1 treated intestinal organoids at 3 days.

(G) Immunoblot analysis of O-GlcNAc (CTD110.6) in intestinal organoids after 3 days of OSMI-1 treatment at increasing concentrations.

(H) Quantification and representative images of clonogenic potential of *Lgr5*$^{eGFP-IRES-CreERT2}$; *Vil*$^{CreER}$ derived GFP$^{hi}$ ISCs treated with vehicle, 20 µM ST045849 or 0.3 µM OSMI-1 treated. N=5. ***p<0.01, ***p<0.001* by one-way ANOVA.

(I) Quantification and representative images of RNAscope *Lgr5*+ cells per human jejunum organoid. Red: *Lgr5*; white: DAPI. N=3, 10 organoids per replicate. *****p<0.0001* by Student's t-test.

**Supplemental figure 3. Decreasing OGN increases PPAR and Lipid programs**

(A) KEGG enrichment of ST045849 treated chondrocytes using GSE275524. Log2FC>1 FDR<0.05.

(B) Overlap of DEGs between ST045849 and GW501516 treated organoids and hypergeometric p value of the overlap.

(C) KEGG enrichment of the genes overlapping between ST045849 and GW501516 (orange), and the genes specific to ST045849 (blue). No enrichment was found in the genes specific to GW501516.

(D) Module score plotting of the genes overlapping between ST045849 and GW501516, the genes specific to ST045849, and the genes specific to GW501516.

(E,F,G) Immunoblot analysis confirming increased abundance of PPAR targets in intestinal organoids treated with vehicle, ST045849, and GW501516 (E), OSMI-1 (F), Azaserine (G).

(H) Quantification and representative images of organoid budding in vehicle and ST045849 3-day, 6-day and 9-day treated intestinal organoids. N=3, 15 organoids per replicate ****p<0.001, ****p<0.0001* by one way ANOVA.

(I) Quantification of lipid droplet area and representative images of MC38 cells stained with BODIPY 493/503 after 3 day ST045849 treatment. N=3, 10 cells per replicate **p<0.05* by Student's t-test. Green: BODIPY; Blue: DAPI.

(J) Representative images of vehicle and ST045849 treated intestinal organoids. Green: PLIN2; Red: Actin; White: DAPI.

**Supplemental figure 4. Decreasing OGN increases PPAR and Lipid programs**

(A) Immunoblot analysis of O-GlcNAC (RL2 clone) and total HistoneH3 as the loading control in ST and vehicle treated *Ppar-d/a*$^{iKO}$ and wild type organoids.

(B) Quantification of clonogenic potential of WT and *Ppar-d/a*$^{iKO}$ organoids after 12 days (3 passages) of ST treatment. N=5, ns: not significant, ***p<0.01* by two-way ANOVA.

(C) Quantification and representative images of clonogenic potential of WT and Cpt1a-deficient *(Cpt1a*$^{iKO}$*: Cpt1a*$^{fl/fl}$*; Vil*$^{CreER}$*; Lgr5*$^{eGFP\text{-}IRES\text{-}CreER}$*)* organoids after 12 days (3 passages) of ST treatment. N=4, ns: not significant, **p<0.05* by two-way ANOVA.

(D) Immunoblot of 3-day treated WT and *Cpt1a*$^{iKO}$ organoids.

(E) Immunoblot of PPAR target proteins from WT, KO(*Ppar-d*$^{iKO}$), +HA-PPARd MC38 cells treated with vehicle or ST045849.

(F) Bar plot of Log2FC expression of *Ogt* in bulkRNAseq of ISCs of iKO's relative to WT in Control diet mice. *FDR<0.05.

(G) Bar plot of Log2FC expression of *Ogt* in bulkRNAseq of ISCs of shown genotypes of HFD relative to Control diet mice. *FDR<0.05.

(H) Box plot of normalized expression of *Ppar-d* in Control and long-term HFD ISCs from GSE151047. Numeric FDR shown on plot.

(I) Log2FC of *Ppar-d* expression in ISCs of HFD and Fasted normalized to Control from GSE67324 and GSE89568 respectively. No significant changes observed.

(J) Immunoblot analysis of MC38 HA-PPARd input, anti-HA, and anti-OGT immunoprecipitated.
