## Supplementary figures and images for "O-GlcNAcylation regulates PPAR-driven metabolic programming in intestinal stem cells"

### Supplemental Figure 1

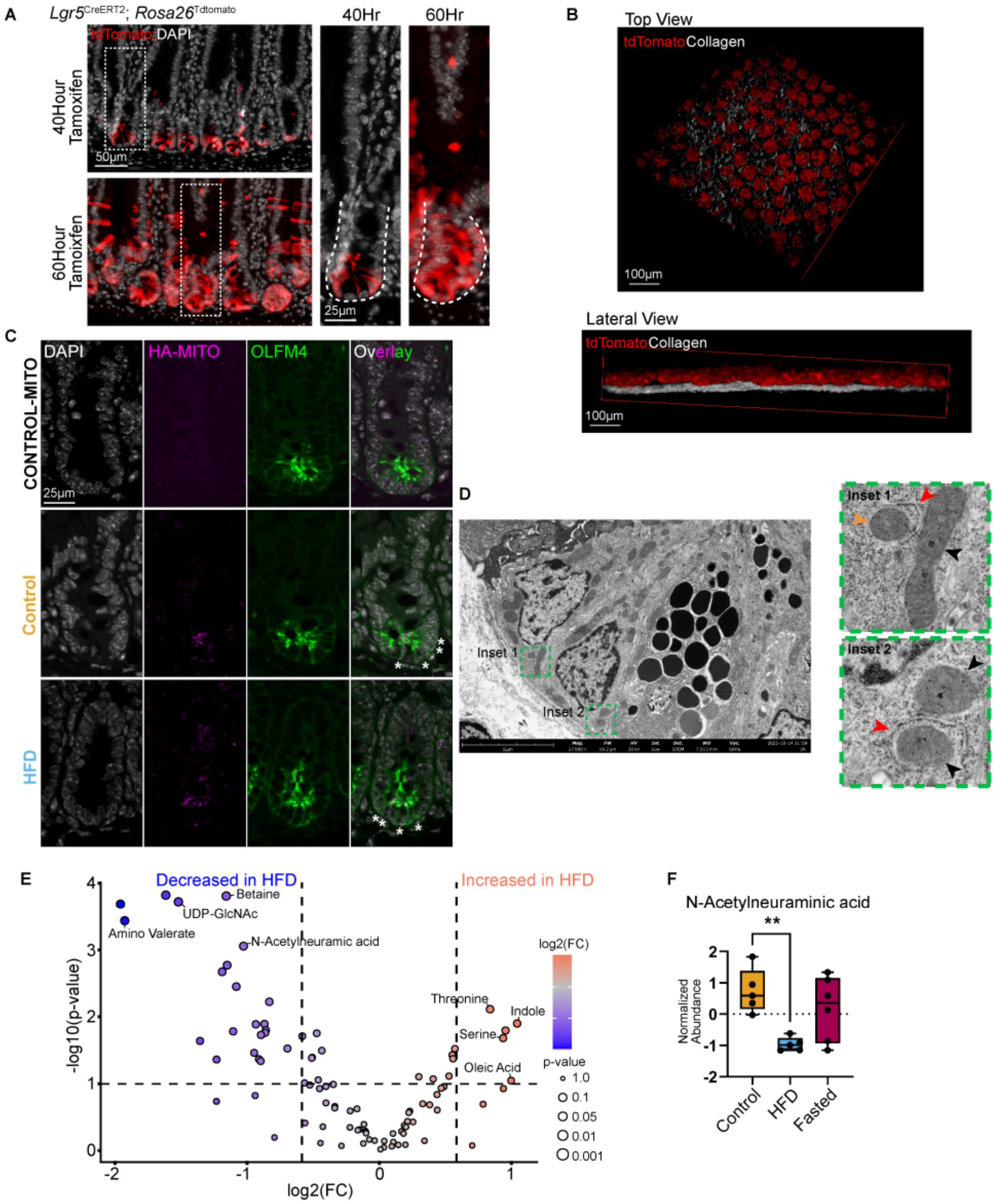

### Supplemental Figure 2

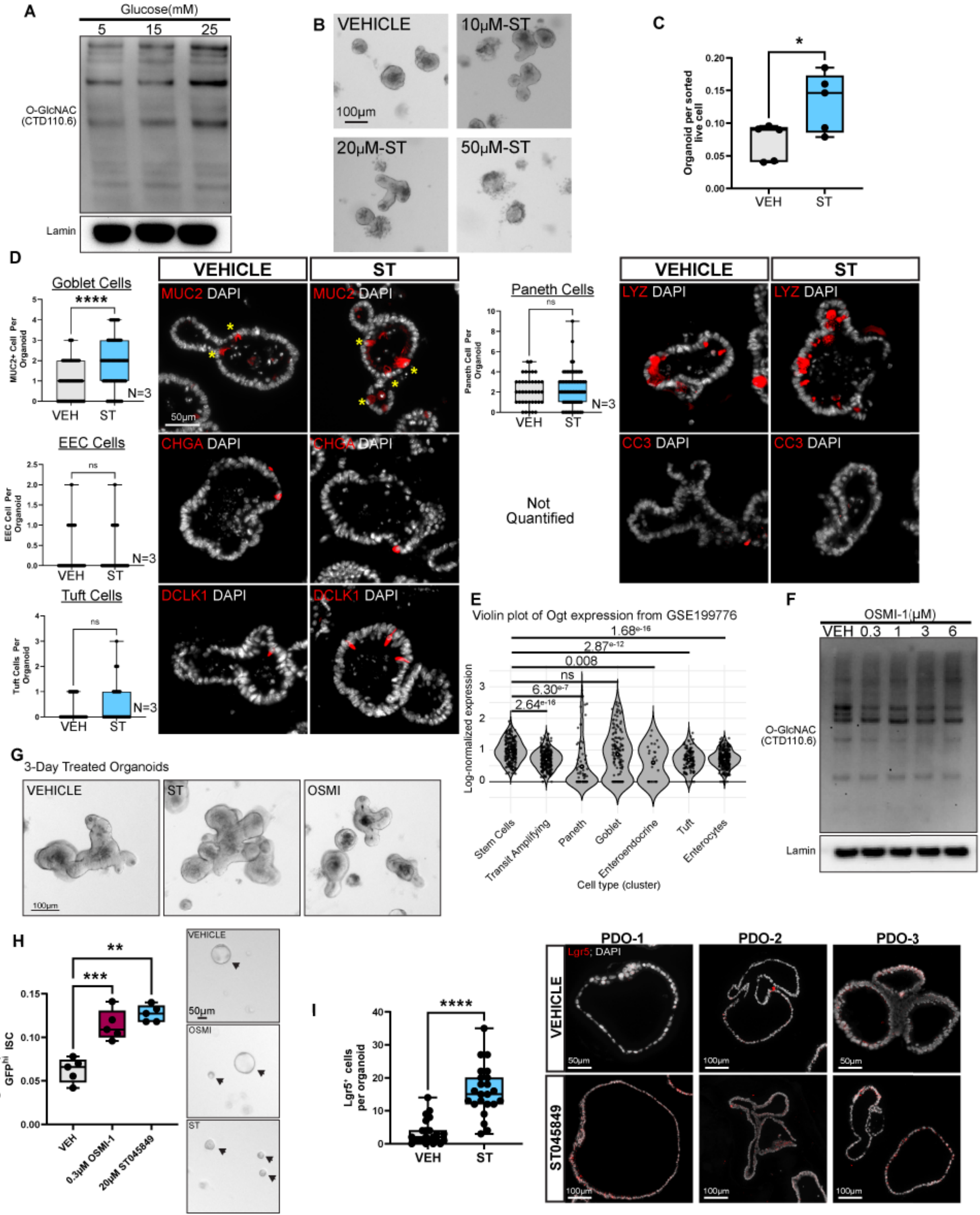

### Supplemental Figure 3

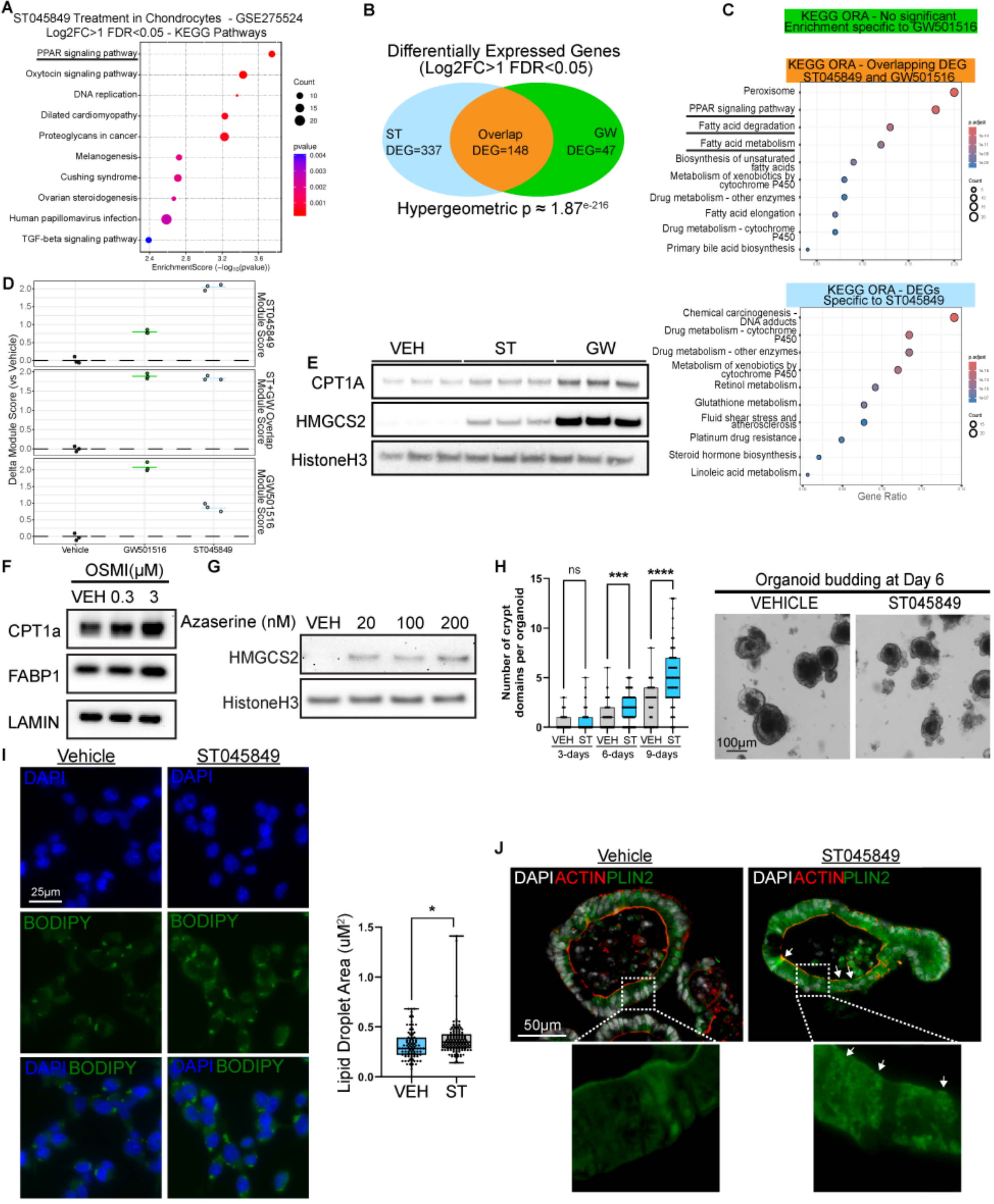

### Supplemental Figure 4

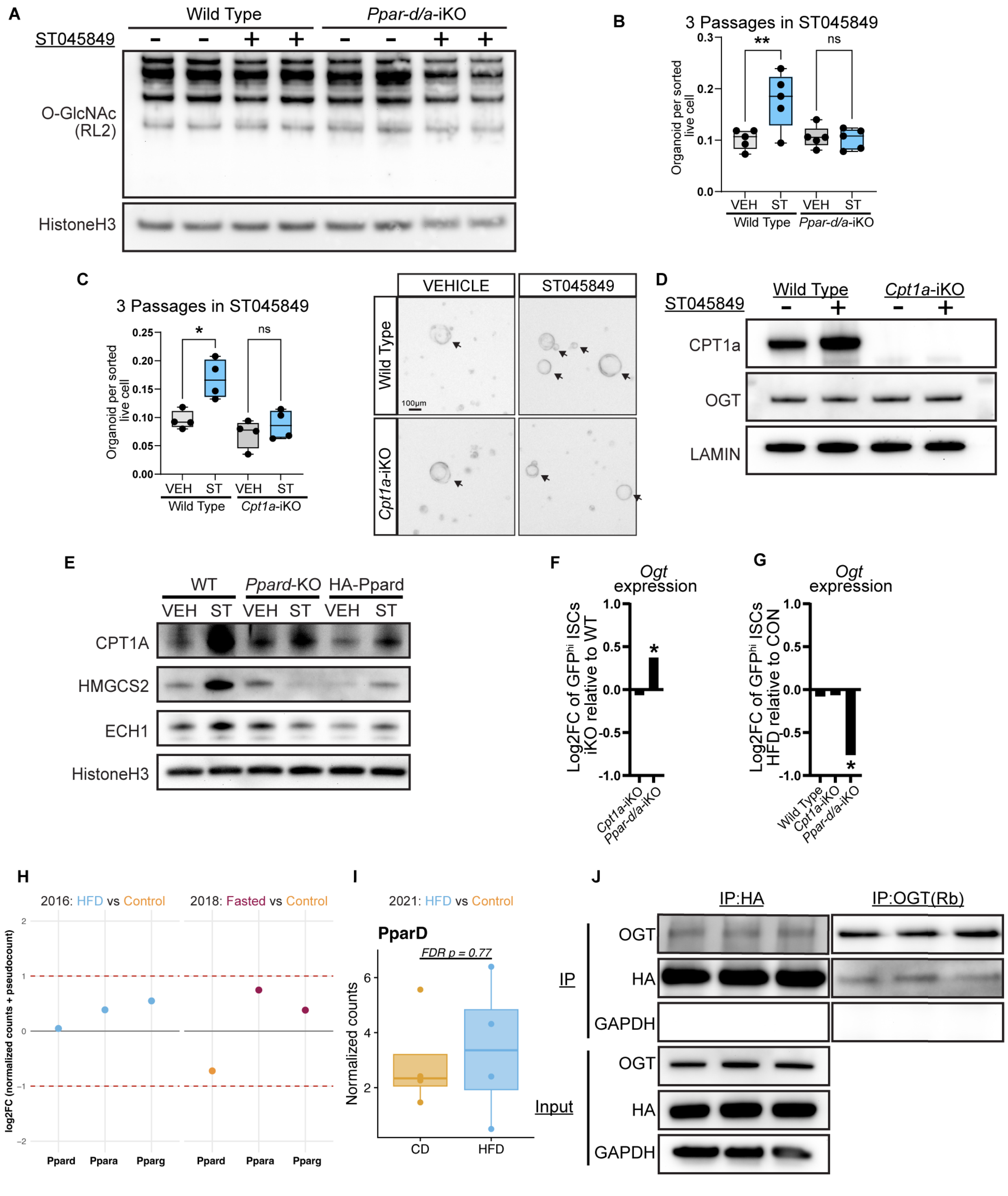
